# Evidence for face selectivity in early vision

**DOI:** 10.1101/2020.03.14.987735

**Authors:** Florence Campana, Jacob G. Martin, Levan Bokeria, Simon Thorpe, Xiong Jiang, Maximilian Riesenhuber

## Abstract

The commonly accepted “simple-to-complex” model of visual processing in the brain posits that visual tasks on complex objects such as faces are based on representations in high-level visual areas. Yet, recent experimental data showing the visual system’s ability to localize faces in natural images within 100ms (Crouzet et al., 2010) challenge the prevalent hierarchical description of the visual system, and instead suggest the hypothesis of face-selectivity in early visual areas. In the present study, we tested this hypothesis with human participants in two eye tracking experiments, an fMRI experiment and an EEG experiment. We found converging evidence for neural representations selective for upright faces in V1/V2, with latencies starting around 40 ms post-stimulus onset. Our findings suggest a revision of the standard “simple-to-complex” model of hierarchical visual processing.

**Significance statement:** Visual processing in the brain is classically described as a series of stages with increasingly complex object representations: early visual areas encode simple visual features (such as oriented bars), and high-level visual areas encode representations for complex objects (such as faces). In the present study, we provide behavioral, fMRI, and EEG evidence for representations of complex objects – namely faces – in early visual areas. Our results challenge the standard “simple-to-complex” model of visual processing, suggesting that it needs to be revised to include neural representations for faces at the lowest levels of the visual hierarchy. Such early object representations would permit the rapid and precise localization of complex objects, as has previously been reported for the object class of faces.

## Introduction

According to the standard model of visual processing (Hubel & Wiesel, 1962; Maunsell & Newsome, 1987; Riesenhuber & Poggio, 2002), simple physical features (e.g., luminance and edges) are encoded first, in low-level visual areas such as V1, while more complex shape features are encoded later in intermediate visual areas such as V4. Complex object categories (e.g., faces and objects) are thought to be encoded in high-level visual areas such as the Fusiform Face Area (FFA).

Humans are able to report the presence of specific objects (e.g. an animal or a vehicle) in natural cluttered scenes as early as 250 ms post-stimulus onset ((Fabre-Thorpe et al., 2003; Poncet et al., 2012; Thorpe et al., 1996; VanRullen and Thorpe, 2001). Those latencies are short, but compatible with a progression of the neural activity from low-level to high-level visual areas, then followed by the formation of a decision-variable in the frontal cortex (Freedman et al., 2003; Riesenhuber & Poggio, 2000). However, the recent use of saccadic response paradigms, rather than manual response paradigms, has revealed that object detection can be achieved substantially faster than previously thought. Faces, for instance, can be saccaded to within 100 ms following stimulus onset (Crouzet et al., 2010). Since conduction delays to generate an oculomotor response take around 20-35 ms (Heeman et al., 2017), this suggests that the visual detection has already been made by 65-80 ms following stimulus onset. Such latencies pose a challenge for the standard hierarchical view of the visual system. A second remarkable feature of these ultra-rapid saccades is their localization accuracy: subjects are able to trigger saccades accurately toward very small faces of 1° visual angle pasted at an eccentricity of 7° in a complex cluttered scene, after only 120 ms (Brilhault et al., 2011). V1/V2 neurons are a possible candidate since their receptive fields subtend 1° in humans at 7° eccentricity (Dumoulin & Wandell, 2008), with the earliest responses starting 45 ms post-stimulus onset (Foxe et al., 2008). By contrast, V4 receptive fields may be too large since they subtend 4°-6° at 6° eccentricity in humans (Motter 2009).

The precision of those saccades in cluttered environments and the fact that they start as early as “express saccades” – the fastest known saccades in humans, that are directed toward luminous dots (Fischer et al., 1984) – suggest that early visual areas, which are characterized by small receptive fields and short activation latencies, might contain representations of complex object categories such as faces.

In the present study, we tested the hypothesis of face-selective representation in early visual areas using a multi-pronged approach. Specifically, to probe the temporal characteristics of putative early face-specific responses, we conducted two eye-tracking experiments and an EEG experiment. To probe the spatial characteristics of putative early face-specific responses, we conducted a functional Magnetic Resonance Imaging (fMRI) experiment. In order to link the results across experiments, stimuli were similar across experimental techniques. They consisted in small faces of 2° size, to match the size of V1/V2 receptive fields. We here report converging evidence that faces elicit specific responses in V1/V2 as early as 40 ms post-stimulus onset. This finding suggests a more nuanced picture of visual hierarchical processing than the traditional “simple-to-complex” model, namely one in which early areas contains neuronal representations selective for complex objects.

## Material and Methods

Across all studies, all subjects were naïve as to the purpose of the study, gave their written informed consent before participating, and were paid for their participation. The Institutional Review Board of Georgetown University approved the study.

### Experiment 1: Specificity of ultra-rapid face detection

#### Experimental Design

##### Participants

Eight (6 women, 22.3 ± 1.9 years old across all subjects) healthy, right-handed human subjects with normal or corrected-to-normal vision took part in the experiment.

##### Stimuli

The stimuli used in the experiment were natural scenes (n=15, resolution 15° × 20° of visual angle, from an online database) in which items (faces and distractors) were pasted. To avoid that pasted items pop-out, the contrast histogram of the pasted item was stretched so that that the bottom 1% and top 1% from the pixels of this item match the bottom 1% and top 1% from the pixel values of the patch of natural scene replaced. All the images were grayscale.

Items were pasted at one of four locations at 7° visual eccentricity from the center of the screen: 25 or 45 ° above and below the horizontal meridian, on the left and right side of the screen. This asymmetrical spatial configuration aimed at activating the primary visual cortex at symmetrical sites on the lower and upper banks of the calcarine fissure (Di Russo et al., 2005) granted that the horizontal meridian is represented in the lower bank of the calcarine fissure (Aine et al., 1996). To approximately match the size of the receptive fields in V1/V2 at 7° eccentricity (Dumoulin & Wandell, 2008), all faces and distractors were resized to span 2° of visual angle (along the height dimension). 7° eccentricity was chosen to be in the range of the eccentricity used in (Brilhault et al., 2011), in which fast saccades toward faces were observed. 15 emotionally neutral faces of males generated with the FaceGen 3.1 software development kit (Singular Inversions, Toronto, Canada) were used. Elements external to the faces (necks and shoulders) were manually removed. From those faces, new face images (inverted, configuration-scrambled and phase-scrambled faces) were created using the Gimp image manipulation program (https://www.gimp.org/) and MATLAB (The Mathworks, MA): “configuration-scrambled” faces in which the internal features (the eyes, the nose and mouth) were positioned at random places, inverted faces, and phase-scrambled face (Xu et al., 2005). 15 images of houses (online database) were also used as distractors (Figure 1A). In this experiment, the “configuration-scrambled” faces, the inverted faces, the phase-scrambled faces and the houses are called “distractors”.

**Figure 1.**
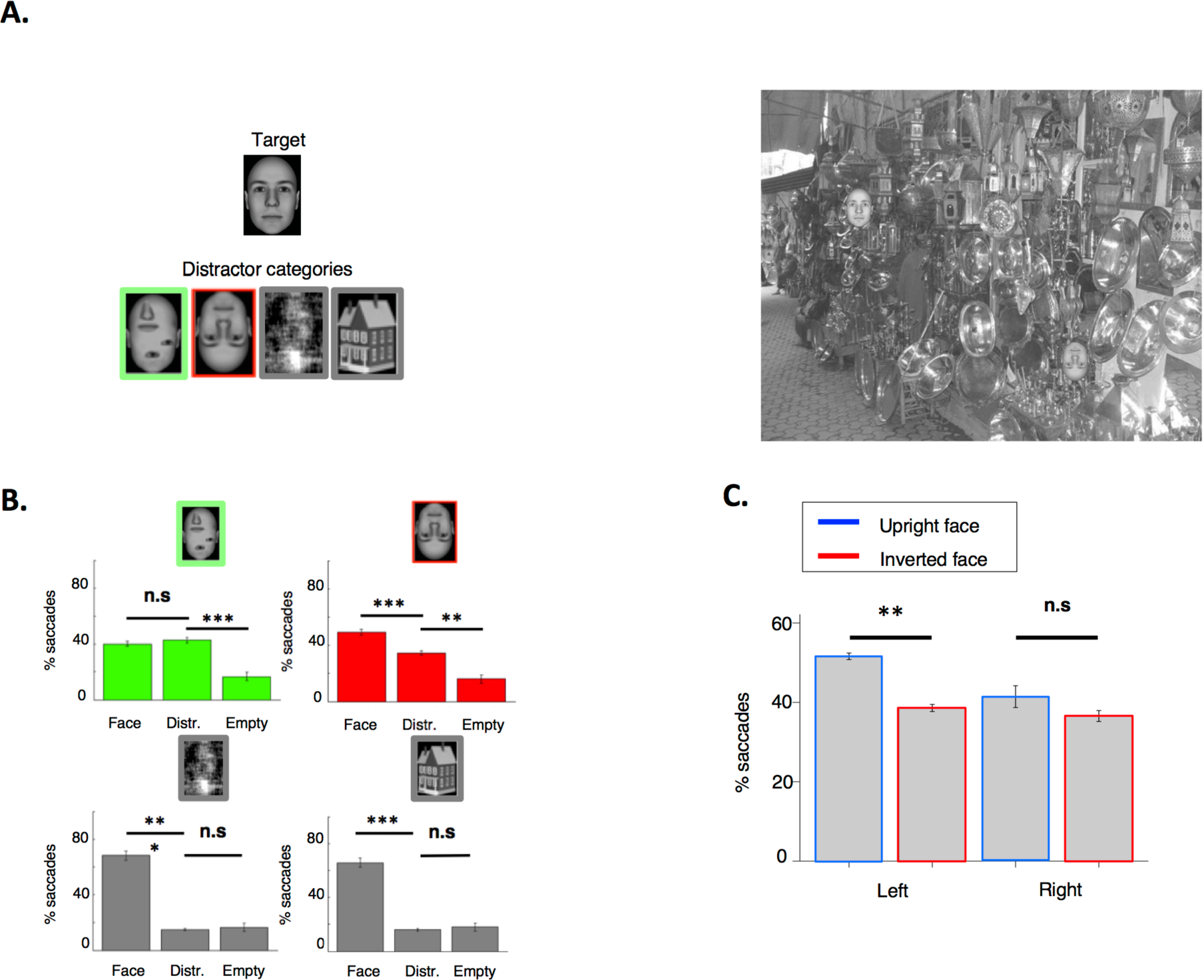
Experiment 1: Fast saccades toward faces have limited selectivity. **A.** Stimuli and paradigm. At each trial, one face and a distractor (from left to right: a configuration-scrambled face, an inverted face, a phase-scrambled face, a house) were displayed at 2 positions, here Left up and Right bottom among 4 positions (Left up, Right up, Left bottom, Right bottom). **B.** Percent of first saccades landing in the quadrant with the face, with the distractor, or toward one of the empty quadrants. Results for configuration-scrambled faces are shown in green, for inverted faces in red, for phase-scrambled faces in gray (left), for houses in gray (right)). In the configuration-scrambled face condition, saccades were oriented as often toward the configuration-scrambled face as toward the target (the upright face). When the distractor was an inverted face, saccades were directed more often toward the target than toward the inverted face distractor. In the control conditions where distractors were houses and phase-scrambled faces, the percent of saccades oriented toward the distractor was similar to the percentage of saccades toward an empty quadrant. **C.** Percent of first saccades in the inverted face condition when the target and the distractor were either both in the left hemifield, or both in the right hemifield. When they were in the left hemifield, saccades were more often directed toward the target than toward the distractor. When they were in the right hemifield, saccades were as often oriented toward the distractor as toward the target. (*** p < 0.001, ** p < 0.005,* p < 0.05). Data are presented as mean ± SEM.

##### Procedure

Stimuli were presented on a computer screen at a viewing distance of 60 cm (refresh rate 60Hz, resolution 1024*768, 24*18° visual angle). Eye movements were recorded using a camera-based eye tracker (SR research, Eyelink 1000 Plus) with a temporal resolution of 2000 Hz. A chin and headrest helped the subjects to stabilize their head. Each trial started with a white fixation cross. Subjects had to keep their eyes on the cross (within a box of radius 1° of visual angle) for 150 ms for the natural scene to be displayed. A 200-ms gray screen was displayed between the end of the fixation and the start of the natural image. This gap enables a faster initiation of saccades (Fischer & Weber, 1993, Kirchner & Thorpe, 2006, Crouzet et al., 2010). Subjects were instructed to saccade as quickly as possible toward the face pasted in the natural scene. The natural scene was displayed until the subject’s eyes gazed at the face for 150 consecutive ms in a square zone centered on the face and of 2° width. Next fixation started after a pseudo-random interval (1000-1200 ms). Subjects performed 15 blocks of 72 trials each. Within a block, there were 48 trials with a face and a distractor (12 possible spatial configurations, one per distractor category), 12 trials with a face only (4 positions, each repeated 3 times), and 12 trials with two faces presented simultaneously (6 possible spatial configurations, each repeated twice). To assess the effect of the different distractor categories independently of face identity, the identity of the face and of the distractor (inverted, scramble or configuration-scrambled) were always the same within an image. Each of the 15 backgrounds and each of the 15 face identities were displayed 72 times (pseudo-random association of a face with a background).

Subjects were informed that, in any trial, the faces would appear at 1 out of 4 possible locations (left up, left bottom, right up, right bottom) (Figure 1A). They were informed that the images would contain distractors, some of them sharing similarities with faces, but that only the “normal faces” were relevant for the task.

#### Statistical Analyses

The position of the left eye from each participant was recorded during the whole experiment, every half ms. Only the first saccade within each trial was kept in the analysis. For each distractor condition, we computed the percentage of saccades landing in the quadrant with a face, in the quadrant with a distractor, and in empty quadrants. This measure was averaged across subjects. We also computed saccadic reaction times, measured as the duration elapsed between the display of the stimulus and the initiation of the saccade, as recorded by the eye tracker. This measure was averaged across subjects too. We computed the minimum saccadic reaction times (SRT), which is a statistical estimate of the latency at which correct saccades start to be more numerous than saccades toward the distractor’s quadrant (Crouzet et al., 2010). For each subject, we computed the cumulative count of number of saccades whose SRT was inferior to a cutoff value, step size 1 ms. We then searched for at least 20 consecutive bins with significantly more correct than erroneous responses using a chi-square test with a criterion of p < 0.05. The first of those bins was considered to correspond to the minimum SRT.

### Experiment 2: Timing and spatial accuracy of ultra-rapid face detection

#### Experimental design

##### Participants

Sixteen (14 women, 23.6 ± 4.1 years old across all subjects) healthy, right-handed human subjects with normal or corrected-to-normal vision took part in the experiment.

##### Stimuli

Stimuli consisted of annuli (radius: 7° of visual angle, width: 2.5 ° of visual angle) cut out from the same database of natural scenes as used in Experiment 1. In each trial, a face (height: 2° of visual angle) was pasted at a random polar angle, always at 7° eccentricity from the center of the screen (Figure 2A). 48 grayscale scenes with a high degree of clutter were selected after visual inspection to make sure that, after the procedure of luminance equalization described above, the pasted faces were still clearly visible at any position within at least 2 out of 4 quadrants in the annulus. 36 grayscale photographs of adult faces of various ethnicities were used (online database) to maximize the ecological value of our results. The neck and shoulders were manually removed from those images as in Experiment 1. Eye movements were recorded using the same setup as in Experiment 1.

**Figure 2.**
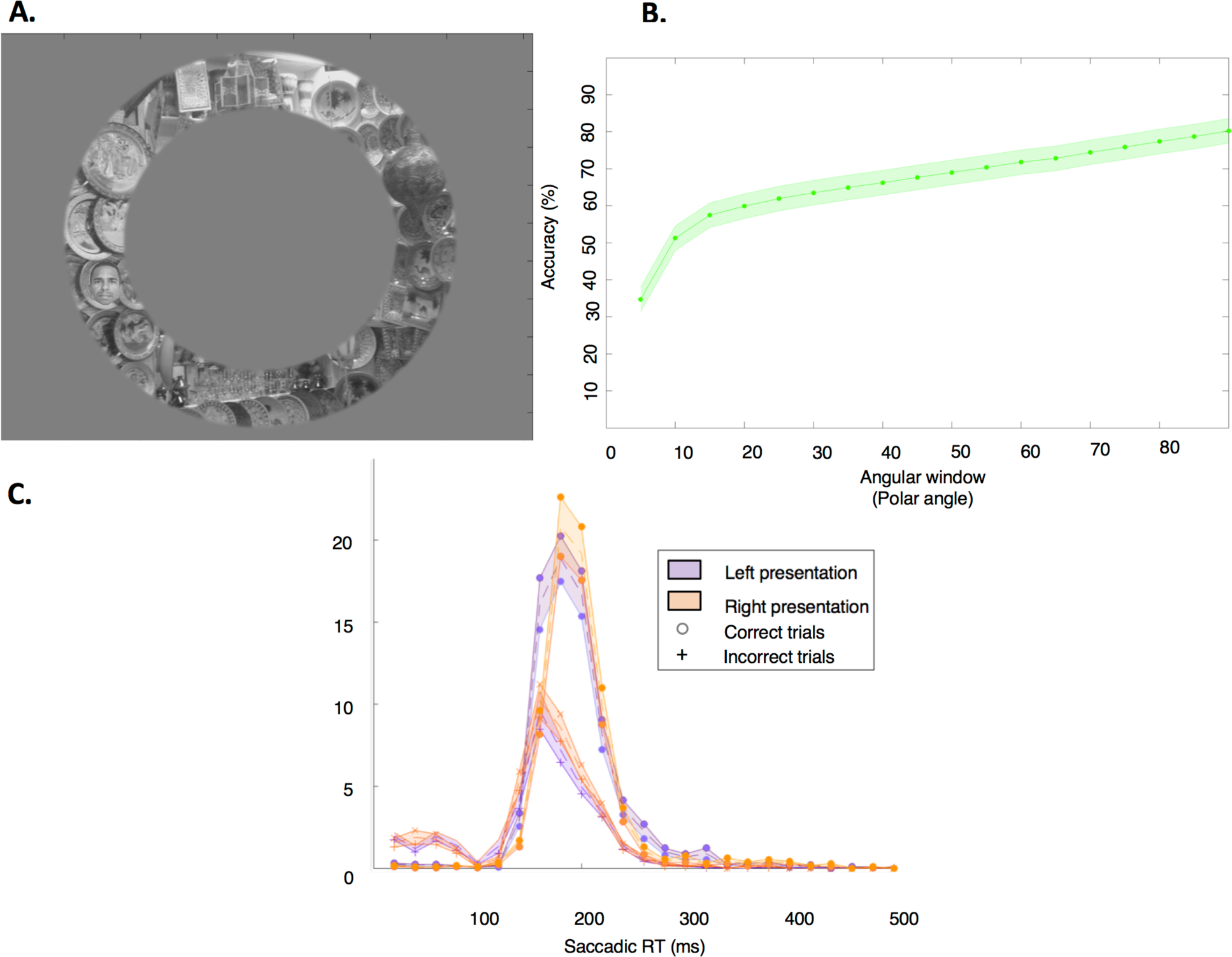
Experiment 2: Faces are saccaded to faster when they are in the left hemifield. **A.** Example stimulus. The face target could be pasted anywhere within an annulized natural scene. **B.** Spatial precision of target saccades in Experiment 2. The plot shows the percentage of saccades landing within an angular window centered on the face, averaged across participants, as a function of the size of the window (degrees of polar angle). The slope rises steeply up to 15 degrees, as progressively more face target-directed saccades are included, then becomes shallower with constant slope, presumably since further increases in window size include progressively more saccades targeted at distractors. The total size of the window is twice the value indicated in abscissa since it extends to both sides. This graph thus reveals that most of the saccades aggregated in an area centered on the face and expanding at ± 15 degrees of polar angle. **C.** Distribution of saccadic reaction times for correct and incorrect saccades, for left-presentation and right-presentation of the faces, averaged across participants. The distribution of correct saccades diverges from the distribution of incorrect saccades earlier for left than for right presentation. This shows that saccades start to be selective earlier for left than for right presentations.

##### Procedure

The experimental setup was identical to that of Experiment 1 except that the distance between the subjects’ eyes and the screen was 93 cm (60 cm for the two first subjects, but stimulus size in degrees of visual angle was identical). 36 faces were displayed, and subjects were instructed to saccade as quickly as possible to the faces. Similarly to Experiment 1, once the participant’s gaze was within the face for 150 consecutive ms, the next trial started after a pseudo-random interval (between 1000 and 1200 ms). Each face was displayed once within each quadrant (36 * 4 = 144 trials).

#### Statistical Analyses

To assess the properties of the earliest saccades that can be selectively oriented toward faces, only the first saccade of each trial was kept for analysis. We computed the minimum saccadic reaction times (SRT) with the same method as described in Experiment 1.

### Experiment 3: EEG

#### Experimental Design

##### Participants

21 healthy (9 women, 24 ± 4.2 years old across old subjects), right-handed human subjects with normal or corrected-to-normal vision took part in the experiment. 2 subjects out of 21 were removed from the group analysis (one since the image was removed in 30.6 % of the trials due to eye movements, the other due to the small amplitude of his N170 upright-face evoked component, at more than 3 std from the mean (average amplitude of the peak, left-presentation group, left-hemisphere: −4.3 ±1 µV, individual value for this participant: −0.49 µV; on the right hemisphere: −4.7 ± 1 µV, individual value: −1.7 µV).

##### Stimuli

100 different grayscale pictures of faces (the same as in the Experiment 2, plus 64 from the same database, from which the neck and shoulders were removed) were used in this experiment.

##### Procedure

The eyetracking setup was identical to that of Experiment 1. In 10 out of the 19 subjects included in the group analysis, eye movements were tracked with the Eyelink SR 1000 (for the other subjects, eye movements could not be recorded because of difficulties with calibrating the eye tracker due to glasses and/or contact reflection). Scalp voltages were measured using an Electrical Geodesics (EGI, Eugene, OR) 128-channel Hydrocel geodesic sensor net and Net Amps 300 amplifier. Incoming data were digitally low-pass filtered at 200 Hz and sampled at 500 Hz using common mode rejection with vertex reference. Impedances were set below 75 kΩ before recording began and maintained below this threshold throughout the recording session with regular impedance checks between blocks.

Half of the subjects (n = 9) were presented with faces in the left visual field, among which 5 had their eye movements recorded. The other half was presented with faces in the right visual field (n = 10), among which 5 had their eye movements recorded. Grayscale pictures of faces were displayed on a gray background on the horizontal meridian. We chose a gray background rather than a natural one to avoid a reduction of the amplitude of face-related signals (Cauchoix et al., 2014). To engage attention to the faces, as in Rossion & Caharel, 2011, we engaged subjects in a face categorization task (upright vs. inverted). The faces were either displayed at 1° eccentricity (height: 0.93° visual angle) or at 7° eccentricity (height: 2° visual angle), with eccentricity chosen randomly for each trial (50 faces at 1° eccentricity, 50 faces at 7° eccentricity). The faces size was determined using the cortical magnification factor in V1 whose value was computed from the formula in (Rousselet et al., 2005) so that, at the two presentation eccentricities, the faces stimulate similar portions of V1. Each trial began with a fixation point displayed for 300–600 ms followed by the brief display of a face, either upright or inverted, for 150 ms. 150ms was chosen as it is longer than our time window of interest (∼50–100 ms) to avoid contamination of possible face-selective signals by stimulus-offset transient responses caused by the stimulus disappearance (Macknik & Livingstone, 1998). Accurate stimulus timing was verified with a photodiode. The subjects then reported whether the face was upright or inverted by pressing the ‘1’ or ‘3’ key, respectively, with their right index and ring fingers (timeout: 2 seconds). After 600 ms, a new trial began. Subjects were asked to keep their eyes on the fixation point during the overall duration of a trial. Subjects were instructed to withhold eye blinks until the end of a trial and could pause the experiment between trials to blink and/or rest. For subjects whose eye movements were recorded, the face appeared only after 150 consecutive ms with the eyes on the fixation spot, thus extending the duration of the fixation in certain trials. For those subjects, the face was removed and the trial aborted if the subject was moving his/her eyes outside of the fixation point during the face presentation. The configuration (upright or inverted), the eccentricity and the identity of faces were pseudo-randomized and counterbalanced across the experiment. Subjects performed 10 blocks of 100 trials each.

#### Statistical Analyses

##### Data Analysis

Data pre-processing and statistical analyses were performed using EEGLAB (Delorme & Makeig, 2004), FieldTrip (Oostenveld et al., 2011) and MATLAB (The Mathworks, MA). Noisy channels were interpolated. Data were band-pass filtered between 0.1 and 30 Hz with a causal filter. A causal filter was used in order to avoid shifts in the latency of the ERPs (Rousselet, 2012). The filtered signal was then epoched from 150 ms before stimulus to 250 ms after the stimulus, baseline corrected in the [-150, 0] ms time window, and trials in which one of the facial channels was exhibiting a variation of more than 75 µV in the [-150, +200] ms time window were removed. Only trials in which the response was correct were kept. Event-related potentials (ERPs) were averaged separately for each subject and for each face orientation (upright, inverted). ERP subject averages were then grand-averaged separately across the subjects with left-face presentation, and across the subjects with right-face presentation. Significant differences between the ERP amplitude in the upright versus inverted conditions were detected (separately within the left-presentation group and the right-presentation groups) using paired t-tests (one-tailed, threshold at 0.05, to test the hypothesis that upright faces elicit a signal of larger amplitude than inverted faces) in a clustering procedure (Oostenveld et al., 2011), with a minimum cluster size threshold of 5 (Scholl et al., 2014). This clustering analysis was performed on all the trials, combining both eccentricity conditions to increase the signal-to-noise ratio. We ran the analysis on the [0-150ms time window], i.e., during the stimulus presentation. Signals were selected over the most posterior channels (n = 51), that broadly overlaid posterior visual areas.

### Experiment 4: fMRI

#### DUMMY

##### Participants

16 healthy, right-handed human subjects (5 women, 20.54 ± 1.1 years old) with normal or corrected-to-normal vision took part in the experiment. Three subjects were excluded from the group analysis: one was singing in the scanner and unable able to focus, the other turned out to be under medication (Truvada), the third one had an excessive amount of head motion (> 8mm).

#### Experimental design

##### Procedure

In the main experiment, stimuli consisted of natural grayscale scenes (the same as in the experiments 1 and 2, 50 different scenes) on which 4 items were pasted (height 2°, at 7° eccentricity, at 25 and 45° above and below the horizontal meridian), to match the location and size of the items in Experiment 1. The items were pasted on gray circles of 4.5° diameter to reduce the effect of variations in the scene content on the BOLD response. Within an image, the four items consisted of an upright item (U), the inverted version of U (I), the scrambled version of U (US), the inverted version of US (IS). U was either a face (50 different photograph of faces, chosen from the same set as in Experiments 2 and 3), or a house (25 different photographs of houses, from an online database) (Figure 4A). The arrangement of the items U, I, IS, US on the four gray circles (top left, top right, bottom left, bottom right) was varied across trials, thereby defining four configurations. The scrambling procedure to create the scrambled items was the one introduced by Stojanoski & Cusack, 2014, which keeps low-level properties of the items: a degree of warping of 35 was used in order to make the face category unrecognizable. Each participant viewed a total of four blocks (125 trials per block, 25 faces * 4 spatial configurations, plus 25 houses). The sets of faces and houses were repeated twice during the experiment, between two blocks, randomized between subjects. Within one block, the 25 house trials had the same spatial configuration, with a different spatial configuration between each block.

##### Main experiment

The purpose of the experimental design in this experiment was to mask the faces and made them invisible to the participants. Each trial (Figure 4B) started with a white fixation cross on a gray background that was displayed for 1000 ms and was followed by a stream of 12 characters pseudo-randomly chosen from a set containing the letters of the alphabet, the numbers 1-9, and 10 punctuation marks ({!, @, #, $, %, ^, &,*, (,)}), and presented in a rapid sequence (5 frames each, 60Hz refresh rate) (adapted from (Beck & Kastner, 2005)). In each trial, a natural scence (with face items or house items, as described above) was embedded in the stream and stayed on the screen for 4 frames. In 70% of the trials, there was an ‘L’ in the stream. Subjects were instructed to press a button as soon as they saw the letter ‘L’. The rapid serial visual presentation aimed at keeping the attention maximally engaged on the center of the screen. The letter ‘L’ was always displayed after the natural scene so that subjects were fully focused on the letter stream at the moment of appearance of the natural scene. The precise timing of the presentation of the stimuli of interest (the natural scene with four faces/four houses) was chosen randomly, it occurred between the 4th to 8th character and could thus not be predicted. Subjects were instructed to keep their gaze and attention on the stream of characters in the middle of the screen. They were told that the natural scenes displayed were irrelevant to the task.

##### MRI acquisition parameters

Data were acquired with a 3T MRI scanner (Magnetom Trio, Siemens) at Georgetown University’s Center for Functional and Molecular Imaging. A 12-channel head coil was used, TR = 2040 ms, TE= 29ms, flip angle = 90°, 35 interleaved axial (thickness = 4.0 mm, no gap; in-plane resolution = 3.2 × 3.2 mm^2^). At the end, 3D T1-weighted MPRAGE images (resolution 1 × 1 × 1 mm^3^) were acquired from each participant. Visual stimuli were back-projected from a computer screen (resolution: 768*1024) on a mirror within the scanner (distance mirror-eyes: 89 cm). Data from four runs of event-related scans, one run of face localizer scan, and two runs of retinotopic localizer scans were acquired from each participant.

##### V1 retinotopic localizer scans

The V1 localizer scans aimed at identifying the portions of V1 (4) activated by presentation of the faces and houses, in order to focus our analyses on those regions. High-contrast achromatic flickering (contrast-reversing at 20Hz) checkerboard patterns were displayed on a gray background (diameter, 3°) successively at the 4 positions occupied by the pasted items (top: left and right; bottom: left and right, at 7° eccentricity). The checkerboards were displayed for 16 consecutive secs at each position and each position was repeated twice. During the presentation of the checkerboards, in the middle of the screen, the letters ‘T’ and ‘L’ were displayed in alternation, in a randomized order (50 ms of presentation per letter, frequency of occurrence of the letter L: 70%). Similar to the main experiment, subjects were instructed to press a button every time they were seeing the letter ‘L’. The first retinotopic localizer scan was run right before the main experiment, the other scan was run right after the main experiment.

##### Face localizer scan

After the second retinotopic localizer scan, a face localizer scan was run to identify the Fusiform Face Area (FFA) in each subject. We used stimuli and a similar design as described in (Kanwisher et al., 1997): 50 different grayscale images of faces and houses, distinct from the ones in the main experiment, were displayed in the center of the screen in blocks on a uniform background. Each block, either with faces or with houses, was run twice. The face and house images were purchased from a commercial source and post-processed using programs written in MATLAB (The Mathworks, MA) to eliminate background and to adjust image size (to 200 * 200 pixels), luminance and contrast. 10 subjects were instructed to passively view the stimuli, 3 others were engaged in a 1-back task to increase attentional engagement.

#### Statistical Analyses

##### Behavioral Data Analysis

We computed the signal detection theoretic (SDT) measures d’ and c (MacMillan & Creelman, 2005) in the target letter detection task. The variable d’ is a measure of a participant stimulus discrimination sensitivity (here, between the letter ‘L’ and other symbols), while c is a measure of a participant bias to report the letter ‘L’. These measures were calculated on the basis of the rate of hits (letter ‘L’ reported in trials in which the letter ‘L’ was present) and false alarms (letter ‘L’ reported in trials in which the letter ‘L’ was absent).

##### MRI Data Analysis

All processing and most statistical analyses were done using the SPM8 software package (http://www.fil.ion.ucl.ac.uk/spm/software/spm8/). The first five volumes of each scan were discarded to allow for scanner equilibration. The functional scans and the localizer scans were preprocessed separately. The echoplanar images (EPIs) were reoriented, then temporally corrected to the middle slice, spatially realigned, and normalized to the Montreal Neurological Institute (MNI) template brain. Images were then smoothed with a full-width at half-maximum of 6 mm Gaussian kernel. After removing low-frequency temporal noise with a high-pass filter (1/128 Hz), fMRI responses were modeled with a design matrix comprising the onset of trial types and movement parameters as regressors using a standard hemodynamic response function.

##### Identification of V1 Regions of interest (ROIs)

V1 regions of interest were identified from the two retinotopic localizer scans, separately for each participant, via the MarsBar toolbox (Brett et al., 2002). Participant-specific ROIs were identified with the contrast of one checkerboard position versus the three other checkerboard positions and this was performed for each of the 4 positions. Each contrast resulted in a focus located in the contralateral hemisphere, below the calcarine sulcus (CS) for checkerboards presented in the upper visual field, and above the CS for checkerboards presented in the lower visual field. We could infer from (Dougherty et al., 2003) that each checkboard (diameter: 3°, at 7° eccentricity) should activate around 75 mm^3^ of cortical surface in V1, i.e. approximately 10 voxels of 8 mm^3^. Based on this estimation, to define V1 ROI, we adjusted the statistical threshold to obtain, in each participant, clusters of approximately 10-15 voxels (significant at the corrected cluster level of at least p < 0.05, else, at the uncorrected cluster level of at least p < 0.001) (Glezer et al., 2015, Jiang et al., 2006). We thus aimed at identifying V1 ROIs, through the use of checkerboard stimuli which are known to strongly activate V1 (Engel et al., 1997), and by restricting the ROIs to 15 voxels. However, for reasons of parsimony, because the activity in V1 propagates to V2, we interpret our data as being in V1/V2, in other words in the early visual cortex.

##### Identification of face-selective ROIs

Face-selective ROIs were identified from the localizer scan separately for each participant, via the MarsBar toolbox (Brett et al., 2002). Epochs with face and house stimuli were modeled with two box-car functions convolved with a canonical hemodynamic response function (HRF). Participant-specific ROIs were defined by voxels that displayed face > house responses. We focused our analysis on the right FFA (Kanwisher et al., 1997, Jiang et al., 2006). ROIs were selected by identifying in each participant the cluster in the right temporal cortex that was significant at the cluster level (corrected cluster level, at least p < 0.05 else, uncorrected cluster level, at least p < 0.001). With such thresholds, only subjects for whom a cluster of at least 30 voxels was found, in a location close to the published location of the right FFA, approximate MNI coordinates (39 ± 3 −40 ± 7 −16 ± 5) (Grill-Spector et al., 2002) were included in the FFA analysis (n = 8 out of 13). In 3 out of the 5 subjects for which no right FFA-cluster could be found with the contrast face > house, we could obtain, through the contrast face > baseline, a right FFA-cluster in a location close to the published location of the right FFA according to Grill-Spector et al., 2002. Analyses were run within the homogenous set of 8 subjects whose FFA was defined via the contrast face > house, and, to increase the statistical power, analyses were also run with the 3 additional subjects with the cluster found via the contrast face > baseline.

## Results

### Experiment 1: Fast saccades toward faces have limited selectivity

We asked subjects to saccade, as fast as they could, toward faces in natural scenes containing a face as well as a distractor sharing, to different degrees, physical properties with faces.

The comparison of the percentage of saccades oriented toward the face versus the distractor revealed that, when the distractor was a configuration-scrambled face (i.e., a face whose internal parts had been scrambled randomly, see Figure 1 A), subjects made as many saccades toward the configuration-scrambled face as toward the target face (Figure 1B), despite the instruction to saccade toward the normal face (% saccades in the quadrant with the normal face: 40.31% ± 1.72%, with a configuration-scrambled face: 42.93% ± 2%, paired t test: T_(7)_ = −1.18, p = 0.27). In trials with an inverted face, a phase-scrambled face or a house, subjects made more saccades toward the face than toward the distractor (inverted face distractors: % saccades in the quadrant with the face: 49.28% ± 2.1%, with an inverted face: 34.63% ± 1.67%, T_(7)_ = 6.6, p = 3 * 10^-4; house distractors: % saccades in the quadrant with the face: 68.35% ± 3.35%, house distractors: 14.9% ± 0.7%, T_(7)_ = 13.89, p = 2 * 10^-6; phase-scrambled distractors: with the face: 65.92% ± 3.53%, with a phase-scrambled face: 16.06% ± 1%, T_(7)_= 11.9, p = 6 * 10^-6). The min SRT, a statistical estimate of the latency at which the first saccades to the (target) face start to be more numerous than toward the distractor (Material and Methods), were, on average, 140 +22 ms, 149 ± 7 ms, 136 ± 11 ms, 138 ± 15 ms (in the configuration-scrambled, inverted face, phase-scrambled, house distractor conditions, resp.). Note that minimum SRTs in this experiment were longer than in Crouzet et al., 2010, presumably since subjects expected the distractors to be similar to the target and therefore potentially adopted a more cautious strategy.

Since saccades to faces have been found to be more accurate and faster when the faces were displayed in the left visual field rather than in the right visual field (Crouzet et al., 2010), we conducted a separate analysis of the saccades when the face and the distractor were both in the left hemifield versus both in the right hemifield. In trials with an inverted face distractor, a higher number of saccades toward the face relative to the distractor was observed for left-hemifield presentation of the face and distractor (% saccades within the quadrant with the face: 51% ± 1%, with an inverted face: 38.6% ± 1%, T_(7)_ = 5.24, p = 0.001). In contrast, for right-hemifield presentation of the face and distractor, subjects made saccades as often toward the inverted face as toward the upright target face (% saccades within the quadrant with the face: 41%, with an inverted face: 36%, paired t test: T_(7)_ = 0.84, p = 0.42) (Figure 1C), suggesting lower face selectivity for right-hemifield presentations. For the other categories, we did not expect such an asymmetry because configuration-scrambled faces act as perfect distractors while houses and phase-scrambled faces do not act as effective distractors.

In summary, Experiment 1 provided evidence that fast face detection has limited shape selectivity, with configuration-scrambled faces frequently being mistaken for targets. This finding is compatible with the notion of fast face detection being based on detectors that encode only face parts, not holistic faces. Supporting this hypothesis, inverted faces, whose parts are affected somewhat by inversion, were not as effective distractors as the configuration-scrambled faces. Another interesting finding from Experiment 1 was a hemispheric asymmetry of fast face detection, with an advantage for face detection in the left visual hemifield. We next conducted another experiment to more finely assess the spatial precision as well as lateralization effects of the earliest saccades oriented toward faces.

### Experiment 2: Faces are saccaded to faster and more precisely when they are in the left hemifield

In Experiment 2, subjects were asked to detect faces in “annulized” natural images consisting of a natural scene windowed by an annulus, which contained a face image (2° of visual angle high). The saccades started as early as 100-120 ms post stimulus onset (Figure 2C). Figure 2B shows that saccades were aggregated in an area centered on the faces and extending up to about 15-20 polar degrees on both sides. We therefore labeled as “correct” the saccades that landed in an area centered on the faces and extended up to 15 degrees both sides and computed the minSRT (Material and Methods). The correct saccades represented 59.6 ± 4% of the total number of saccades and landed at 1.07° ± 0.05° visual angle from the center of the face, with an average latency of 182.3 ms (average of the moment of onset of the first saccades). The minimum SRT (Material and Methods) was shorter for left presentations than for right presentations (minimal SRT, on average, left-presentation: 160 ± 4 ms, right presentation: 177 ± 4 ms, paired t-test, T_(15)_ = −3.2, p = 0.008) (Figure 2C). Furthermore, saccades landed closer to the face in left-presentation trials relatively to right-presentation trials (left presentation: 3.54 ° ± 0.28 °, right presentation: 4.06 ° ± 0.25 °, paired t-test: T_(15)_ = −2.8, p = 0.013).

### Experiment 3: EEG data reveal face-specific responses 40 to 110 ms post-stimulus onset for left-hemifield presentations

To probe the neural bases of fast face localization, we conducted a high-density electroencephalography (EEG) study and assessed the earliest latency of face-selective neural signals. We presented faces in the periphery while asking subjects to look at a central fixation spot and perform an upright/inverted face classification task on peripherally presented faces. They were presented on one side of the screen only, with presentation side alternating between subjects. Akin to previous studies of higher-level face processing (Kanwisher et al., 1998; Rossion et al., 2000), and motivated by the observation of an advantage for upright vs. inverted faces in fast face detection (Experiment 1), we adopted a stringent test to establish face selectivity of neural responses, requiring significantly different neural responses to upright vs. inverted faces. This contrast avoids confounds arising from low-level stimulus differences between faces and comparison objects (e.g., when comparing responses to faces versus houses, which can be differentiated by other low-level features such as different luminance distributions). To engage attention to the faces, as in (Rossion & Caharel, 2011), we engaged subjects in a face categorization task (upright vs. inverted).

Reaction times in the upright/inverted face categorization tasks were computed on the trials kept for the EEG analysis, i.e., on the correct trials. Across the 10 subjects whose eye movements were recorded, 11.7 ± 2.8 % of the trials on average were aborted due to eye movements. Subjects were faster responding to upright faces than inverted faces (paired t-test, mean average reaction time for correct trials, left-presentation group, upright faces: 538 ± 28.3 ms, inverted faces: 586 ± 35 ms, T_(8)_ = −3.6, p = 0.0069; right-presentation group, upright faces: 603 ± 48 ms, inverted faces: 651 ± 48.5 ms, T_(9)_ = −4.4, p = 0.0016; note that these reaction times were substantially longer than the 0-100ms time window of interest). Reaction times were not significantly different between the left- and right-presentation groups (unpaired t-test: T_(18)_ = −1.05, p = 0.3). Accuracy was very high, and similar between the groups (unpaired t-test, mean accuracy, left-presentation group: 93.3 ± 1%, right-presentation group: 91.3 ± 2.5%, T_(18)_ = 0.72, p = 0.47). The average number of trials kept for the EEG analysis was similar in the left- and in the right-presentation groups (unpaired t-test, left-presentation group: 705 ± 43 trials, right-presentation group: 625 ± 27 trials, T_(18)_ = 1.6, p > 0.1).

A clustering analysis was done using Fieldtrip (Oostenveld et al., 2011) between 0-150 ms relatively to stimulus onset over the most posterior channels, which capture (early) visual-evoked responses and notably V1 activity (Foxe & Simpson, 2002). In the left-presentation group, a significant cluster (p = 0.035) differentiating the signal in the upright and inverted conditions between 40 and 110 ms was found (Figure 3A). The cluster was lateralized in the right hemisphere (Figure 3B). In the right-presentation group, the cluster started later, extending from 98 to 140 ms (p = 0.04) (Figure 3C). The cluster was initially lateralized in the left hemisphere (Figure 3D). A control analysis was run to verify that the early clusters with a significantly higher signal amplitude for upright vs. inverted faces were not driven by subjects without eye recordings, who could have moved their eyes despite the instruction not to do so. Specifically, we computed the mean signal amplitude for each condition (upright and inverted configurations) within the subjects whose eye movements were recorded. We predicted that for participants whose eye movements were recorded and who maintained fixation throughout each trial (Material and Methods), thus with no possible contamination of the ERP by eye movements, we should observe a significant difference between the upright vs. inverted conditions in the ERP. In this analysis, we combined the participants with left-hemifield and right-hemifield presentation in order to increase statistical power. This analysis revealed that the differences were indeed observed within those subjects (group of subjects with eye movements recorded, within the appropriate cluster according to the presentation side, upright condition: 1.16 ± 0.36, inverted condition: 0.87 ± 0.32, T_(10)_ = 3.9, p = 0.0029). This confirms that the early face selectivity could not be explained by differential movements toward upright versus inverted faces.

**Figure 3.**
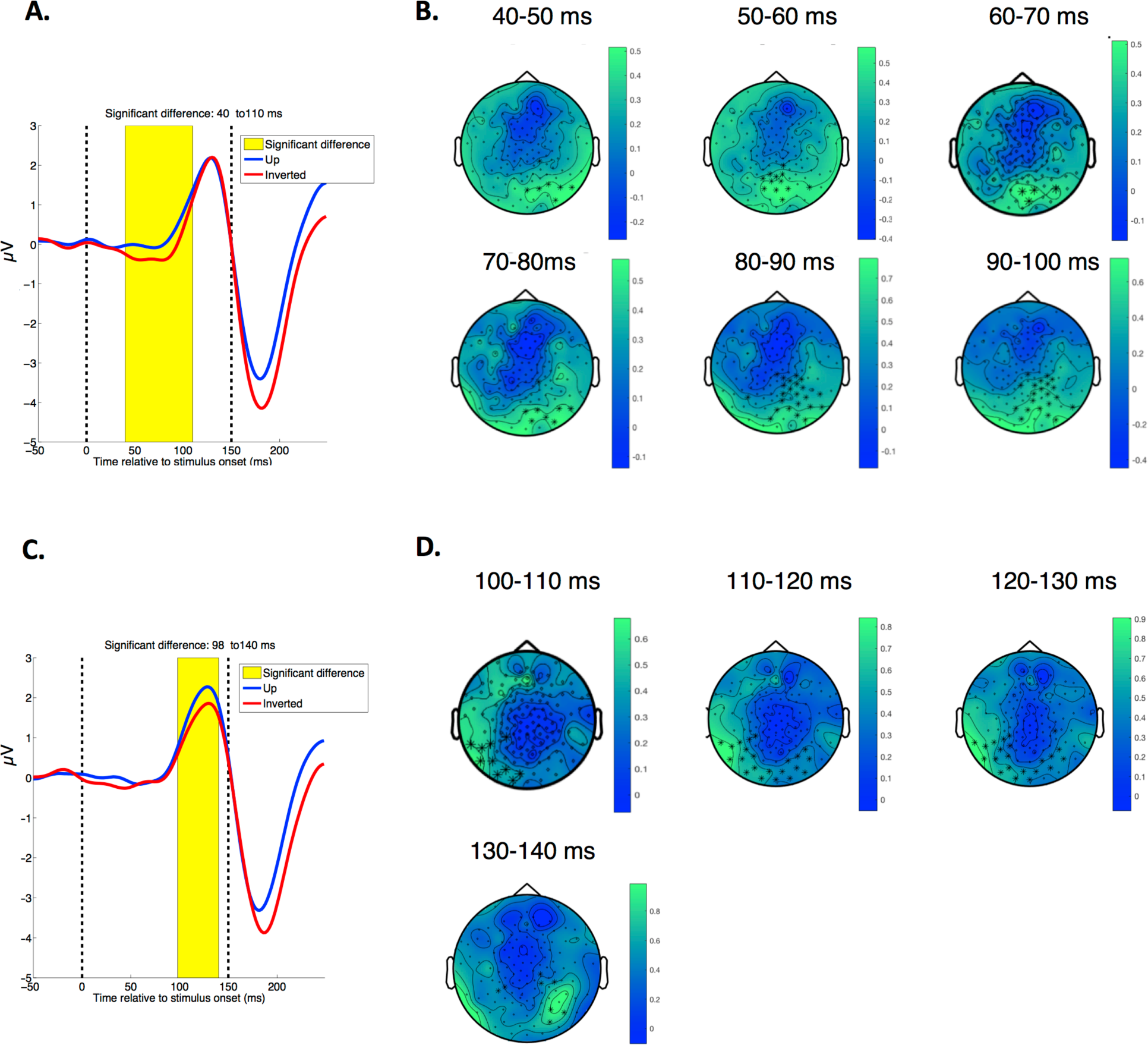
Experiment 3: EEG data reveal significant face-specific responses starting at 40 ms post-stimulus onset for left-stimulus presentations. Event-related potentials (ERPs) computed within the cluster found in the time window [0-150] ms. **Left-presentation group. A.** Mean ERPs elicited by upright versus inverted faces, computed across the channels belonging to the cluster. The cluster ranges between [40-110] ms post-stimulus. The time window over which the clustering analysis was run is indicated by dashed lines, the window of significance is in yellow. **B.** Time course of the cluster, electrodes significant during the whole time window (10 ms bins) are indicated by stars: the cluster is initially lateralized in the right hemisphere. **Right-presentation group. C.** Mean ERPs elicited by upright versus inverted faces, computed across the channels belonging to the cluster. The cluster ranges between [98-140] ms post-stimulus **D.** Time course of the cluster, electrodes significant during the whole time window (10 ms bins) are indicated by stars: the cluster is initially lateralized in the left hemisphere. It becomes more spread-out starting around 110-120 ms poststimulus onset.

**Figure 4.**
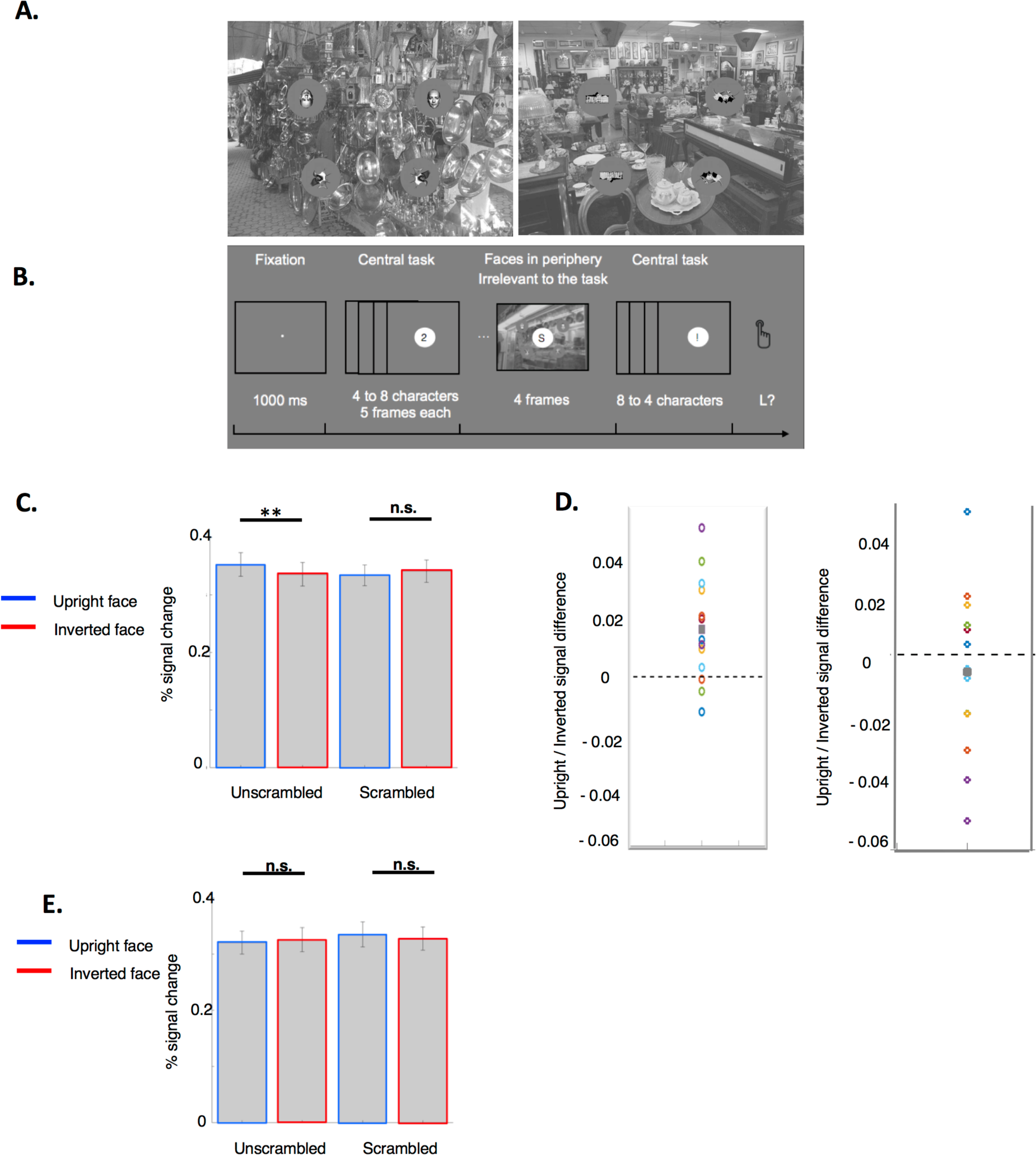
Experiment 4: fMRI data provide evidence for face-specific responses in V1/V2. **A.** Example stimuli. Left: example of a face trial, one face is pasted in its 4 different versions (inverted, upright, scrambled, and inverted scrambled on the top left, top right, bottom left and bottom right respectively) in the middle of gray circles. Right: example of a house-trial, built in the same way as the face-trial. **B**. Experimental design. Participants performed a central letter detection task in a rapid visual serial stream. In this stream, a natural image with faces (or houses) at 7° eccentricity was displayed at an unpredictable moment. Subjects were not told about the presence of faces/houses. **C.** Percent signal change within V1/V2 ROIs averaged within stimulus category across the ROIs activated by left visual presentations (top left and bottom left). There is a significantly higher percent signal change in trials with upright (unscrambled) faces relatively to trials with inverted (unscrambled) faces, but no significant difference for the comparison upright (scrambled) faces versus inverted (scrambled faces). **D**. Left. Representation of the difference between the percent signal change evoked by the upright (unscrambled) faces and by the inverted (unscrambled) faces, for left-hemifield presentations. Right. Representation of the difference between the percent signal change evoked by the upright (scrambled) faces and by the inverted (scrambled) faces, for left-hemifield presentations. Each dot corresponds to one participant. The average value of those points is represented by a gray, filled, square. **E**. Percent signal change averaged within stimulus category across the ROIs activated by right visual presentations (top right and bottom right). The percent signal change elicited by upright and inverted faces in this visual hemifield is not significantly different, both for unscrambled and scrambled faces.

### Experiment 4: fMRI data provide evidence for face-specific responses in V1/V2

To identify the sources of the early face-selective neural signal, we conducted an fMRI study. Subjects were engaged in a central letter detection task during a rapid serial visual presentation (RSVP) of symbols and letters, while small faces and houses (2°) were displayed in the periphery (7°). In the central letter detection task, subjects had, on average, a high d-prime that stayed stable across blocks, thus suggesting that they stayed engaged over the whole duration of the experiment (across the four blocks, mean, d’ = 3.3, mean criterion c: −0.8; repeated-measures ANOVA over criterion c with the single factor Block, F_(1, 12)_ = 2.05 p = 0.12). Debriefing at the end of the experiment revealed that none of the subjects noticed either faces or houses items.

#### V1-response to the face stimuli from different categories

We used upright faces (U), inverted faces (I), scrambled upright (US) and scrambled inverted (IS) faces in order to probe for face selectivity in V1. Similarly to the EEG Experiment (Experiment 3), we used inverted faces to assess face selectivity. We also used scrambled faces as a control. The scrambling procedure (from Stojanoski & Cusack, 2014, described in the Methods section) preserved the faces’ basic visual properties, to which the earliest stages of the visual processing are sensitive, while making faces unrecognizable as faces. We reasoned as follows: if there is some selectivity to faces in V1, the BOLD signal in response to U faces should be higher than for the other three categories. The contrast {US vs. IS} enabled us to check that a differential BOLD signal between U versus I items does not arise from an inversion effect but is truly a response specific to upright faces.

Retinotopic localizer scans with flickering checkerboards known to evoke strong responses in V1 (Engel et al., 1997) were used to define the regions of V1 activated during the functional runs by our visual stimulation in periphery (4 items pasted at 7° eccentricity, see Figure 4A). The four ROIs were in the lower and upper bank of the left and right calcarine sulcus, corresponding to the upper and lower presentation of the checkerboards respectively (Material and Methods, mean coordinates of the ROIs: activated by top left presentations: (12 ± 1 −78 ± 1 −5 ± 1), bottom left: (12 ± 1.1 −92 ± 1 21 ± 2), top right: (−10 ± 1 −81 ± 1.3 −8 ± 1.2), bottom right: (−8 ± 1 −97 ± 1 16 ± 2)). The ROI subtended approximately 15 voxels (mean size of the ROI: 13.9 ± 0.68, activated by top left presentations: 13.07 ± 0.67, bottom left presentations: 12.08 ± 0.53, top right presentations: 14.2 ± 0.94, bottom right presentations: 16.15 ± 2.5, repeated-measures ANOVA over the ROIs size, factor ROI location: F_(1,12)_ = 0.13, p = 0.25).

We first considered the percent signal change collapsed across the four ROIs. Responses in V1/V2 to upright and inverted faces were similar, regardless of scrambling status: the unscrambled faces (mean % signal change, U: 0.34 ± 0.04, I: 0.33 ± 0.04, paired t-test, T_(12)_ = 1.6, p = 0.13), and the scrambled faces (mean % signal change, US: 0.33 ± 0.04, IS: 0.33 ± 0.04, paired t-test, T_(12)_ = −0.0026, p = 0.99). A repeated-measures ANOVA on percent signal change showed that the amplitude of the percent signal change in the V1 ROIs was similar between the scrambled and the unscrambled pairs (main effect, factor Orientation (upright/inverted): F_(1, 12)_ = 0.028, p = 0.87, factor Scrambling (scrambled/unscrambled): F_(1, 12)_ = 0.69, p = 0.42, interaction Orientation × Scrambling: F_(1, 12)_ = 0.87, p = 0.37).

Next, based on the lateralization effects found in behavior and EEG, we analyzed the BOLD contrast responses by hemifield (mean size of the two ROIs from left presentations: 12.69 ± 0.61, of the two ROIs from right presentations: 15.19 ± 1.9, paired t-test, T_(12)_ = −1.7, p = 0.1). Interestingly, in the right hemisphere ROIs (activated by left hemifield presentations), the percent signal change in response to upright faces was significantly higher than to inverted faces for unscrambled faces (mean % signal change, U: 0.35 ± 0.036, I: 0.34 ± 0.04, paired t-test, T_(12)_ = 3.19, p = 0.0077 (Figure 4C), with 10 out of 13 subjects with a higher percent signal change for the upright face trials) (Figure 4D, left). In contrast, for scrambled faces, responses to upright versus inverted stimuli was similar (mean % signal change, US: 0.33 ± 0.036, IS: 0.34 ± 0.039, paired t-test, T_(12)_ = −0.8, p = 0.43, with 6 out of 13 subjects with a higher percent signal change for the upright scrambled face trials (Figure 4D, right)). There also was no significant difference between the response to I and IS (T_(12)_ = 0.63, p = 0.53), nor between the response to I and US, as predicted (T_(12)_ = 0.17, p = 0.86). By contrast, when the analysis was restricted to the left hemisphere ROIs (activated by right hemifield presentations), no significant difference between the percent signal change to upright versus inverted faces was found in V1/V2, neither for the unscrambled faces (mean % signal change, U: 0.32 ± 0.041, I: 0.32 ± 0.043, paired t-test, T_(12)_ = −0.75, p = 0.46), nor for the scrambled faces (mean % signal change, US: 0.33 ± 0.045, IS: 0.32 ± 0.044, paired t-test, T_(12)_ = 0.55, p = 0.58) (Figure 4E). We performed a repeated-measures ANOVA on percent signal change, with factors Orientation and Scrambling to check whether there was a difference in the BOLD-signal for U vs. I faces that was specific to the unscrambled faces. This ANOVA, when performed for left-hemifield presentation, confirmed that the response to upright versus inverted faces was different between the scrambled and the unscrambled conditions (main effect, factor Orientation (upright/inverted): F_(1, 12)_ = 0.6, p = 0.43, factor Scrambling (scrambled/unscrambled): F_(1, 12)_ = 0.97, p = 0.34, interaction Orientation × Scrambling: F_(1, 12)_ = 4.36, p = 0.05). By contrast, for right-hemifield presentation, the response to upright versus inverted faces was similar between the scrambled and the unscrambled conditions (main effect, factor Orientation (upright/inverted): F_(1, 12)_ = 1.18, p = 0.29, factor Scrambling (scrambled/unscrambled): F_(1, 12)_ = 0.014 p = 0.9, interaction Orientation × Scrambling: F_(1, 12)_ = 0.75, p = 0.41). This difference between the left versus right presentation paralleled the findings in behavior (Experiments 1 and 2) and EEG (Experiment 3).

To avoid saccades, we flashed the stimuli very briefly (67 ms), a time too short for eye movements to the stimuli of interest since express saccades, the fastest oriented-saccades in humans, peak at 100 ms (Fischer & Ramsperger, 1984). Furthermore, the debriefing showed that subjects were unaware of the presence of faces in the trials. It is therefore unlikely that the difference in the percent signal change between trials with upright and inverted faces (left hemifield presentations, unscrambled faces) reflected differential patterns in eye movements. Additionally, if eye movements toward peripheral items drove this effect, we would expect that the higher the effect size in the BOLD signal was, the worse the accuracy should have been in the central task. Yet, no correlation was found between the effect size in the BOLD signal (computed as the contrast in percent signal change to I vs. U, analysis restricted to left hemifield presentations) and behavior, both when considering the d’-measure (Pearson correlation, r = 0.34, p = 0.25), or when considering the criterion (Pearson correlation, r = 0.32, p = 0.28).

The FFA is known to exhibit a larger BOLD signal in response to faces than to houses (Liu et al., 2002). Therefore, we performed a control analysis in the FFA ROIs in order to check whether the differential response in V1/V2 ROIs to upright vs. inverted faces presentations could be due to a differential response to upright versus inverted faces in the FFA, that would feed back to early visual areas. The right FFA could be identified in 8 participants through the contrast face > house (mean coordinates: (43 ± 1.41 −49 ± 2.44 −20 ± 2.07), mean size: 103.3 ± 22.81 voxels). Interestingly, while, in the localizer scans, the FFA ROIs were identified via the higher BOLD-signal for face presentations relatively to house presentations, in the functional scans that included the flashed peripheral presentation of small faces and houses (Material and Methods for details), no significant difference was observed in the percent signal change between face and house presentation trials (mean percent signal change in face trials: 0.27 ± 0.025, in house trials: 0.27 ± 0.034, paired t-test, T_(7)_ = 0.13, p = 0.89). There was no significant difference either between the percent signal change in face trials split by the hemifield of presentation of the unscrambled faces, to which the FFA is usually responsive (mean percent signal change, unscrambled faces on the left: 0.26 ± 0.027 on the right: 0.28 ± 0.02, paired t-test, T_(7)_ = −0.84 p = 0.42). To test the robustness of these null effects, we ran the same analysis while adding three more participants (n = 11 out of 13) for which an FFA-cluster could be identified when using the contrast face > baseline (across the 11 participants, mean Talairach coordinates: (40.9 ± 1.58 −52.7 ± 3.41 −19.45 ± 1.77), mean size: 93.9 ± 17.21 voxels). Adding those three participants did not change the results – no significant difference was found in trials with faces versus houses (mean percent signal change, in face trials: 0.27 ± 0.018, in house trials: 0.28 ± 0.028, paired t-test, T_(10)_ = −0.38, p = 0.7), and no difference between trials with faces on the left versus on the right hemifield (mean percent signal change, unscrambled faces on the left: 0.27 ± 0.02 on the right: 0.28 ± 0.013, paired t-test, T_(10)_ = −0.68 p = 0.5). Thus, in our experiment, the small, peripherally presented faces did not elicit face-specific activation in the FFA.

## Discussion

In the present study, motivated by the remarkably short latency and high localization accuracy of fast saccades to faces, we investigated the hypothesis that the early visual cortex contains selective neural representations for complex objects of high ecological importance, specifically faces. Experiment 1 showed that, compatible with a location in lower visual areas, the specificity of fast saccades to faces is moderate, with “configuration-scrambled” and inverted faces interfering substantially with the localization of upright faces. Experiments 1 and 2 also revealed a key signature of fast saccades to faces, namely an asymmetry between the two hemifields that is characterized by a higher selectivity for upright vs. inverted faces for left hemifield presentations (Experiment 1), spatially more accurate and faster saccades for this hemifield (Experiment 2). Second, an EEG experiment (Experiment 3) revealed that event-related potentials (ERPs) differentiated between upright and inverted faces in the left hemifield from 40 ms post-stimulus onset, i.e., in the latency of the first estimated responses in V1 as measured on the scalp (Foxe et al., 2008). For faces displayed in the right visual hemifield, the earliest significant difference in the ERPs between upright and inverted faces started later, at 98 ms post-stimulus onset. This asymmetry between the hemifields in the EEG results therefore matched the asymmetry found in the behavioral experiments. It is worth noting that the EEG difference in latency of the early face-selective response between the left-presentation and right-presentation groups was larger than the difference in minimum saccadic reaction times in Experiment 2. However, participants in that experiment were not only slower for right hemifield presentations but also less accurate. This speed-accuracy tradeoff suggests that behavioral latency differences did not fully compensate neuronal latency differences in Experiment 2. Third, an fMRI experiment (Experiment 4) revealed that in V1/V2, upright faces elicited a higher percent signal change in the BOLD signal than inverted faces. Again, this face-specific response was found specifically for faces displayed in the left visual hemifield, but not for faces displayed in the right visual hemifield. Thus, we found a high degree of agreement between the behavioral, EEG, and fMRI results suggestive of face-selective neuronal populations in early visual cortex.

Our results come in the wake of recent reports suggestive of face selectivity in early vision. EEG classification results indicate that EEG potentials recorded over occipital locations can predict the location of faces as early as 50ms post-stimulus onset (Martin et al., 2014). In an MEG study, the authors reported, for left (but notably not right) presentations, larger responses to face-like vs. house-like patterns in early visual areas with short latencies (Shigihara & Zeki, 2014), but the comparison of responses across different channel groupings for different stimuli in that study left open the specificity of the observed effect. A recent ECoG study (Matsuzaki et al., 2015) reported response differences between upright and inverted faces within 40-90ms following stimulus onset over early visual areas V1 and V2. However, most of the electrodes (80%) were placed in locations corresponding to the upper visual field, therefore resulting in different physical stimulation in the upright face trials versus the inverted face trials. Thus, the differential response might have been elicited by a difference in the physical stimulation. In Uyar et al., 2015, the authors reported a differential response to faces vs. houses in early visual areas. However, it is problematic to use the face vs. house contrast as a marker for face selectivity in V1 since a differential response to faces versus houses can be driven by low-level differences between the two stimulus classes, given the known selectivity of V1 neurons for low-level features (e.g. luminance, spatial frequency). For this reason, to provide a more selective probe for face selectivity, our study used inverted faces, which have been used as a specific control for face processing in a large number of studies (Haxby et al., 1999; Itier & Taylor, 2002; Kanwisher et al., 1998). As a further test, we created scrambled faces that are unrecognizable as faces yet preserve the original faces’ low-level properties; we found that unscrambled inverted faces and scrambled faces in their upright and inverted version elicited a similar percent BOLD signal change in early visual areas ROIs, lower than the signal elicited by unscrambled upright faces. This strongly confirms that the response in the early visual cortex that we found is a response specific to faces as an abstract category, that cannot be accounted for by low-level properties of faces.

The left / right asymmetry that we found across the experiments parallels the well-known left hemifield advantage found for face recognition and categorization (Gilbert & Bakan, 1973, Rhodes, 1985, Sergent & Bindra, 1981) and the right hemisphere advantage for neural representation of faces (Yovel et al., 2008). It has been postulated that asymmetries in visual processing between the left and right hemisphere relate to the level of processing (local vs. global), spatial frequency content of stimuli, or the visual pathway (magno- vs. parvocellular) (for a review, see (Hellige, 1996)). While the mechanisms responsible for the face processing asymmetry are beyond the scope of this paper, it might be of significant interest for future studies to directly investigate the relationship between the low-level face processing asymmetry with the asymmetry of high-level face processing (e.g., in the FFA).

### A new view of hierarchical processing in the visual system

Our study suggests the intriguing hypothesis that there might be neural representations selective for certain complex object classes (such as faces) at the lowest levels of the visual hierarchy. This finding challenges the standard simple-to-complex hierarchical description of the visual system according to which complex object categories are processed at high levels of the visual hierarchy. It is unlikely that the selectivity in early visual areas for upright faces identified in our study (Experiment 4) is the result of feedback activity from areas belonging to the core network for face perception, such as the FFA or the OFA (occipital face area) (Haxby et al., 2000). First, in our study, the usual contrast of Faces vs. Houses, to which the FFA is normally responsive (Liu et al., 2002), did not reveal any significant difference of the percent BOLD signal. This might be due to the fact that the faces were presented in the periphery while the FFA is mostly responsive to central face presentations (Levy et al., 2001). Second, the debriefing conducted at the end of Experiment 4 indicated that subjects did not notice the presence of faces or houses. Those subjective reports suggest that the masking was efficient, thus preventing feedback-driven face-specific activity in V1/V2 (Fahrenfort et al., 2007). Taken together, the early latency of the face-specific response found in the EEG signals and the localization of the face-specific response in V1/V2 suggest that face-specific responses emerge during a first forward sweep of processing and truly reflect a selectivity to faces in early vision rather than the propagation in feedback of the activity from higher-level areas.

Given its position at a low level of the visual processing hierarchy, we would not expect that the degree of face selectivity in V1/V2 would be comparable to the high degree of face selectivity in the FFA, which has been shown to be involved in the discrimination of different faces identities (for a review, see Kanwisher et al., 2006). In fact, the observation that the fast saccades were attracted toward the configuration-scrambled faces (Experiment 1) suggests that face representations in V1/V2 are not selective for the configuration of the faces, in line with findings about face representations in higher areas of occipital cortex that are not selective for the face-part configurations (the Occipital Face Area, OFA) (Pitcher et al., 2007; Liu et al 2010), in contrast to face representations further downstream in fusiform cortex (Andrews et al., 2010; Liu et al., 2010).

The face-specific neural response that we observed in V1/V2, starting at 40 ms post-stimulus onset, constitutes a likely underpinning for the fast saccadic detection of faces. Such a model of “shortcuts” in hierarchical visual processing, with early visual representations directly providing input to decision circuits for fast motor responses is compatible with recent theories of thalamocortical processing (Sherman, 2016) in which corticofugal projections from layer 5 in early visual areas could carry task-relevant signals directly to the superior colliculus. It will be very interesting in future work to explore the plasticity of this circuit, to see whether it is possible to learn fast saccades for new object classes in addition to faces by effecting object-specific plasticity in early visual areas.

## Author contribution

F.C, M.R., X.J., J.G.M., S.J.T designed experiments. F.C. ran experiments with assistance from L.B. F.C. and M.R. analyzed the data. All co-authors contributed to the writing of the manuscript.

## Acknowledgments

This research was supported by National Eye Institute Grant R01EY024161 and a grant from the Agence Nationale de la Recherche ANR-13-NEUC-004-01. We thank Dr. Alex Todorov for face stimuli, and Drs. Bobby Stojanoski and Rhodri Cusack for scrambling code, and Rebekah Farris for help with experiments.

